# The Software Crisis of Synthetic Biology

**DOI:** 10.1101/041640

**Authors:** Sergi Valverde, Manuel Porcar, Juli Peretó, Ricard V. Solé

## Abstract

In fifteen years, Synthetic Biology (SB) has moved from proof-of-concept designs to several flagship achievements. Standardisation efforts are still under way, basic engineering concepts such as modularity and orthogonality are still controversial in biology, and making predictions from computer models is still unreliable. A deep characterization in the pattern of re-use of biological blocks in SB has not been attempted to date. We have compared the topological organisation of two different technological networks, one associated to a standard, large-scale software repository and the second provided by the Registry of Standard Biological Parts (RSBP). Our results strongly suggest that software engineering, and not industrial engineering, is the closest complex system to SB. In both cases, combining standard or quasi-standard components assembly with tinkering may not be at odds with success.

## I. INTRODUCTION

Any technology experiences a phase of explosive growth as a consequence of the combinatorial potential that pervades innovation (Arthur 2009). As more inventions become available, the repertoire of possible artefacts that can be created using previous pieces, designs or modules expands exponentially. The textbook example is provided by electronics (Williams 1985). Starting from the first large, clumsy and inefficient transistor, a rise of Information Technology (IT) became a reality due to the open-ended nature of combinatorial logic (any circuit can be designed from small pieces) and the parallel development of software engineering (Mens and Demeyer 2008). Actually, these two domains coevolved over time, completely changing our relationship with information and design. Synthetic Biology (SB) is an emerging field that offers a similar potential to change the biotechnology landscape. Largely inspired in standard IT, it focuses on the translation into the biological realm of engineering pillars such as modularity, orthogonality and standarization (Purnick and Weiss 2009).

In just one decade, SB has moved from proof-of-concept designs to several flagship achievements attributed to SB-driven strategies. These include microbial drug synthesis, production of new biofuels or novel approaches to disease treatment (Church et al., 2014). The newborn discipline has risen great expectations but also considerable hype. One particular claim is the capacity of SB to exploit combinatorial design principles as the engine of novel, more complex living machines by assembling functionally self-contained parts such as promoters, coding sequences, ribosome binding sites, terminators or protein domains. In this context, the informational nature of living systems (Maynard Smith 2000, Nurse 2008) supports an analogous path of technological development. But the LEGO-like notion of the field has been seriously questioned (Kwok 2010) thus raising two questions. The first is the value of the similarities between IT and SB. The second, whether the landscape of SB designs is growing as a consequence of combinatorial design. In order to answer these questions, a quantitative, systems-view of the technological landscape is needed.

## II. HARDWARE OR SOFTWARE?

Mounting concerns have emerged in relation to the ambiguity of the industrial engineering and electronic metaphoric pools (de Lorenzo, 2011; Porcar et al., 2015), on the difficulties for living things to fit rational design (Collins et al., 2014) or on the inexactitude of the very concept of cells as biomachines (Nicholson, 2013). Are designed constructs like small, widely re-usable components? (Andrianantoandro et al., 2006). If not, its potential as a fast expanding, successful technology might be questionable. What kind of approach can be used to prove or disprove the appropriateness of the analogy? Because the currently adopted representation of cells and synthetic circuits is in terms of logic gates, genetic parts might appear closer to hardware components. In this context, it has been recognised that genetic pieces are actively read-out as a source of algorithmic instructions (Walker and Davies 2013). By contrast, transcription factors (and other parts of the molecular cell machinery) work as hardware operating on genetic instructions. If we need to compare technological fields, a new comparative frame could be software engineering. Using system-level, network approaches, it has been shown that software systems can be described as complex webs of interacting parts (Valverde et al., 2002; Myers, 2003; Potanin et al., 2005) characterised by an extensive reuse of subsystems. More importantly, there are remarkable convergent traits shared between large software structures and molecular cell networks (Solé et al., 2011). In this paper, we will identify global patterns of interaction in networks of co-occurrence of genetic parts used in SB designs in order to search for a fingerprint of combinatorial reuse.

There is an additional reason to consider software engineering here. A specially relevant connection between software development and the potential future of synthetic biology stems from the serious challenges experienced by the former shortly after it started to rise. Already in the 1960s, rising concerns emerged once software projects started to become more complicated, poorly specified and prone to unexpected failures and defects. The failure of software to match continuous advances in hardware constrained the capacity of programmers to effectively use machine characteristics (Dijkstra, 1972). The main cause of this failure was a combination of poor quality and the lack of reusability of software components. Both factors led to chronic maintenance problems (Hooper and Chester 1991). A proposal to solve this “software crisis” was based on the reuse of interchangeable components (McIlroy, 1976). This reuse-based approach to software development, although promising, had limited applicability except for libraries of highly-specialised domains, e.g., mathematical routines. Instead of a successful standardisation, a multiplicity of libraries, codes and programming languages and specialised domains emerged. Soon it became clear that even reliable parts, once in a new context, are not necessarily reliable (Coulange 1998). Some of the problems were solved two decades later while others still persist today (Garlan et al., 1995; Mili et al., 1995). For most practitioners of SB design, the previous story might sound familiar.

## III. GENETIC PARTS VERSUS NETWORKS OF PARTS

The most used and well characterised compilation of biological parts are BioBricks*™*, a repository of up to 7822 DNA sequences, most of which issued from the international Genetically Engineered Machine (iGEM) competition. Critical meta-analyses on the performance of Biobricks are scarce (Vilanova and Porcar, 2014). In the present work, we have tackled for the first time a global analysis of the biological components from the Registry of Standard Biological parts (http://parts.igem.org/Registry_API), and compared its structure and re-use features with both LEGO© (Crompton, 2012) and a large software system (Valverde et al., 2002).

The LEGO© metaphor has been often used within SB (Cserer and Seiringer, 2009; Collins, 2012) as a way of referring to the combinatorial potential: not surprisingly, the maximum award in iGEM is given in the form of an aluminium BioBrick trophy. There is a statistical pattern, which is shared by all the systems discussed here. It is given by the rank-frequency distribution, which gives the number of times *N(r)* that the r-th most common element has been found in a given collection of Lego sets, displayed in Fig. 1a (Crompton, 2012). In Fig. 1b we plot (blue dots) the corresponding distribution for the frequency of use of genetic parts. A similar pattern is shared by the frequency of use of software components within the JAVA library, a good example of object-oriented software (Fig. 1b, red dots). Here too, most elements in the library are used in a very specific way, whereas a few are widely used. In both cases, *N(r)* follows the so called Zipf’s law (Newman, 2005) for large *r*. Zipf’s law is characterised by a power law decay where frequency decays inversely with rank, i. e. *N(r)* ~ r^-γ^ with γ=1. The origins of this fat-tailed statistics has been studied and are known to be a consequence of extensive reuse of existing structures by developers (Valverde and Solé, 2005).

**FIG. 1.**
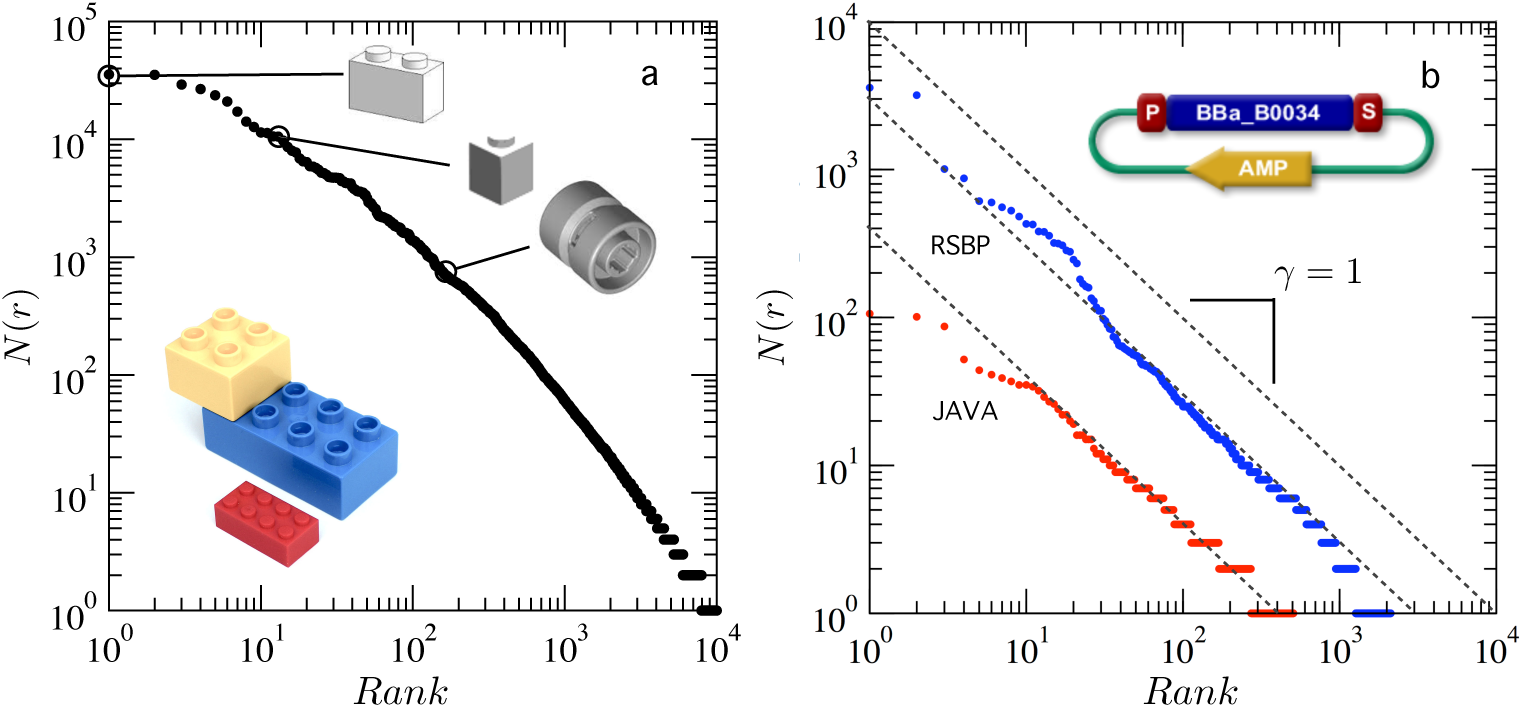
Frequency use of LEGO, software and genetic parts. Three systems are represented: (a) LEGO pieces, (b) genetic parts from the Registry of Standard Biological Parts (RSBP, blue) and software components within the JAVA library (red). Here we display, using a log-log scale, the rank-frequency distributions, where the most frequent element has rank *r* = 1, followed by the next more frequent *r* = 2 and so on. The *r-th* point gives the frequency *N(r)* of use of the *r*-ranked piece. The first set (a) gives the frequency of block use in Lego constructions (Crompton 2012) and we indicate the exact pieces for *r* = 1,10,100. More “generalist” pieces have lower ranks and widespread use. The other two distributions (b) also indicate how often a given part (biological or technological) participates in any engineered design (a genetic construct or a package). Both have long tails that fit Zipf’s law, i e. *N(r) ~ r*^-γ^ with γ = 1, as indicated by the dashed lines.

Both distributions are highly skewed, indicating that they are dominated by a few, very common pieces while most pieces are rare. This indicates that a handful of objects are easily incorporated (the “generalists”, i. e. small pieces capable of gluing together other elements) while the majority are specialised. These are broad statistical distributions that can be obtained from different types of mechanisms. Having fat-tailed distributions is no guarantee that reuse is the leading mechanism. To test this possibility, we need to measure the actual interactions among parts.

## IV. NETWORK STRUCTURE AND PATTERNS OF REUSE

The frequency-rank distribution does not incorporate the real architecture of technological complexity, which is associated to the way elements connect to each other. This means that, in order to get a global picture of the organisation of these systems, a network approach is required (Dogorovtsev and Mendes 2003; Newman, 2010; Valverde and Solé, 2005). By considering the links existing among gates within electronic circuits (Ferrer et al., 2001), libraries within software projects (Valverde et al., 2002; Myers, 2003; Potanin et al., 2005), gene networks (Teichmann and Babu, 2004) or influences among inventions (Valverde et al., 2007), it is possible to unravel the origins of the networks connecting logic components, programs or patents. Of note, several studies have shown that genetic networks and software programs share common structural principles (Yan et al., 2010). To further investigate the parallelisms and differences between software and BioBricks, we performed a network analysis of dependencies among JAVA components and genetic parts.

From the Registry of Standard Biological Parts (RSBP) we reconstructed the graph (hereafter *G*_*R*_) of logical dependencies among parts. The way this was done is illustrated in Fig. 2a with an example. Here, a node represents a biological part and a link between two parts indicates that they are simultaneously engineered within the same construct. The arrows point from a given object to its constituents. The result of this is the BioBrick network, which is composed by several disconnected subgraphs. Most parts in the Registry (88%) belong to one single, large subgraph (Fig. 2b) which we have studied in detail. A few highly connected components create a dense central region, while the majority of nodes are, in average, associated with a small number of other parts. For comparison, in Fig. 2c we also display the largest connected component of the JAVA software network (hereafter *G*_*R*_). In both cases, links describe the use dependencies among either software (Valverde et al., 2002) or biological parts. The different shape of both graphs suggests very different global patterns of organisation, as confirmed by our quantitative study. One obvious and significant difference is the distinct density of connections. The JAVA graph is sparse, as most technological networks, exhibiting a reduced average number of links (average degree) resulting from a supervised pruning of redundant relationships. This is one desirable feature that is far from present in the RSBP graph, where many connections seem to indicate that no such supervised process is at work. Instead, as new parts are added, they are likely to be connected to other parts irrespective of the increasing redundancies that can arise.

**FIG. 2.**
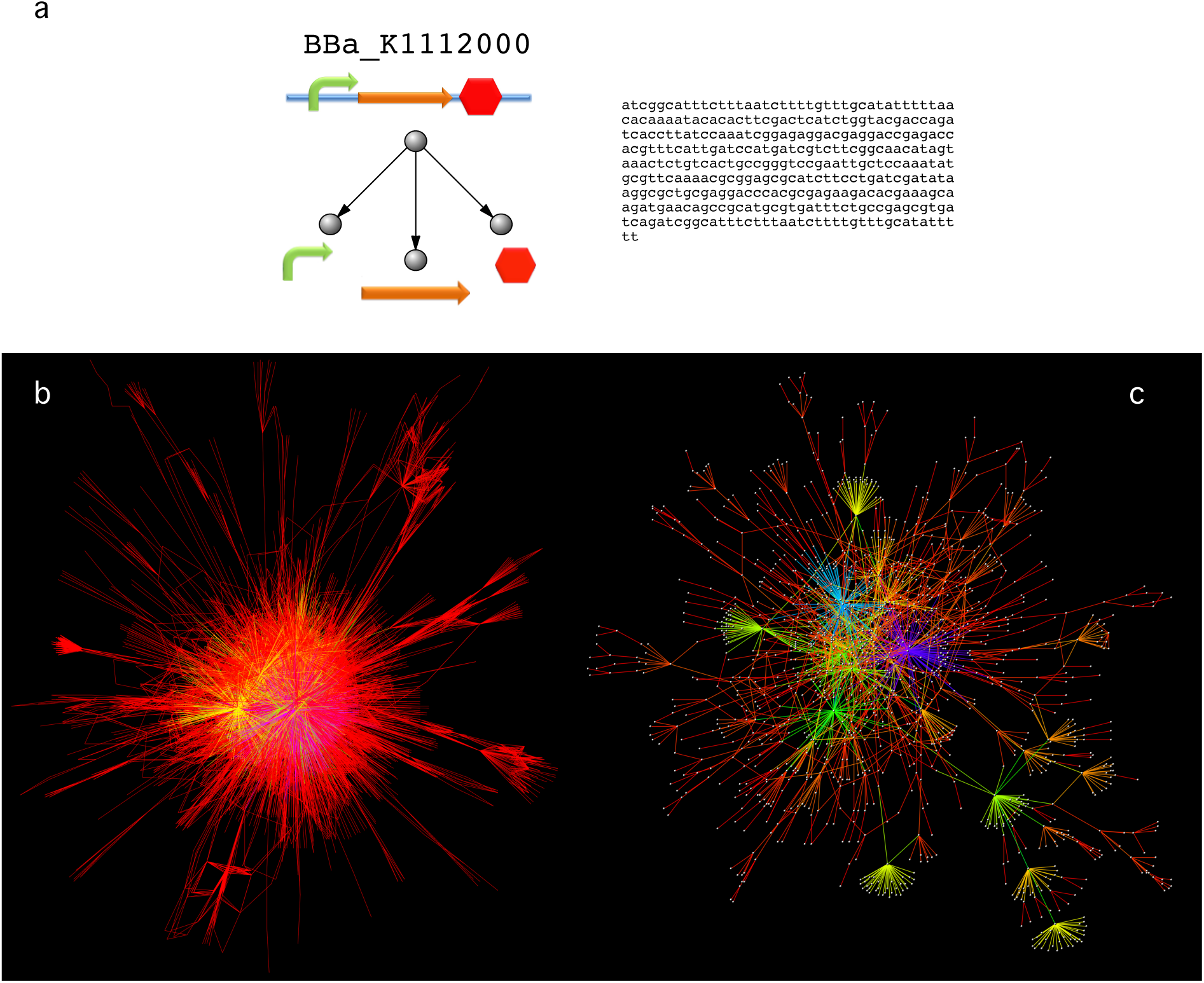
The landscape of software networks versus synthetic biology webs of genetic parts. (a) A network can be defined using genetic parts as components: any two parts that have been used to build a new construct are connected by a link. The global pattern of links is shown in (b) for the Biobrick network and, (c) for the JAVA network of software dependencies.

A common trait in many complex networks is the presence of *Small-World* (SW) effect (Watts and Strogatz, 1998). It is characterised by two key traits: (a) very small path lengths d_*L*_, i. e. short paths separating any two arbitrary nodes; and (b) a high clustering coefficient *C:* many triangles are present, indicating that two nodes sharing a given node are likely to be also connected between them. This feature has proved in systems with extensive reuse. The standard definition of the SW assumes two basic conditions: (1) *d*_L_ ≈ *d*_*rand*_ and (2) *C ≫ C*_*rand*_; in which we compare the values of *d*_L_ and *C* with the expected values *d*_*rand*_ and *C*_*rand*_ of a random version of the networks with all links randomly assigned. Technological webs such as *G*_*J*_ are small worlds: here we have *d*_L_*(G_J_)* = 5.4, *d*_*rand*_(*G*_*J*_) = 6.28 and *C(G*_*J*_) = 0.016 whereas the Registry graph is clearly not a SW because *d*_L_(*G*_*R*_) = 3.42 and *C(G*_*R*_) = 0.0002. Here, we have a very low clustering (close to random) and the path length is below the random expectation *d_rand_(G*_*R*_) = 4.14.

Dramatic differences arise when looking at the modular structure of these webs. A desirable, universal feature of both biological and technological networks is the presence of modularity (Baldwin and Clark, 2000). Modular patterns provide a source of robust local division of labor within a complex system: damage or malfunction of parts within a module do not propagate outside it. In a network context, a module is (roughly speaking) a subsystem such that its components are more connected among them than with the rest of the network. Modularity is measured by means of a parameter *Q* that weights to what extent a given network exhibits modular architecture (Newman, 2010). Here Q is close to 1 if the system is composed of very well-defined, weakly disconnected modules; whereas low values indicate that the system is weakly (or not) modular. Standard systems as JAVA have high *Q* values (here *Q*_*J*_ = 0.75) revealing that the technology is characterised by modules of functionally related components but also displaying multiple dependencies between these modules. By contrast, the Registry network has a rather small *Q* value (*Q*_*R*_ = 0.3), very far from its theoretical maximum. Such a lack of modular architecture tells us that there are no communities of more or less functionally related pieces. Almost everything connects with the “trivial” hubs, with little more organisation. Other measurements confirm this picture. Taken together, our results indicate that the Registry graph is far from a well-organised modular network, and thus lacking the expected attributes of a technological system with differentiated subparts.

## V. LESSONS FOR THE FUTURE

The comparative analysis we report here between biological parts and software components reveals that the landscape of SB, as captured by the network of dependencies, is completely different from the scenario predicted by extensive reuse. Our results reveal that the current iGEM repository strongly deviates from the desirable scenario where different parts are easily combined with others. Instead, it simply shows that a few parts are very often used as almost inevitable components with no additional reuse. The immediate consequence of this is a very limited potential for innovative combinations. As it happened with the early crisis of software development, the promise of reusability within SB will not be met unless new directions are taken.

Although the poor reusability potential of the iGEM landscape paints a rather grim picture, we might gain some insight from the historical development of software. In the early years of IT, programming required to get very close to hardware: programming involved so-called machine code, essentially writing chains of 0’s and 1’s and it was difficult to write and very time-consuming. The first language, FORTRAN, was easy to learn and simplified software development. At that time, it was assumed that the programming language would eliminate coding and debugging, solving problems at a low cost. Just ten years later, these expectations completely failed to be fulfilled. It was discovered that bugs are always present and considerable effort is invested in finding and correcting them (McIlroy, 1976). Poor programming practices and inappropriate integration in large software projects have led to disaster (Hey and Pśpay, 2015): in 2002, software bugs cost the US economy an estimated $59 billion annually, or about 0.6 percent of the gross domestic product (Tassey, 2002). Correcting and maintaining large software projects cannot be achieved without a cost and good solutions have only been obtained after the introduction of software engineering. Nevertheless, we can claim an overall success for software as a major technology.

As for the SB engineering practices, two main points should be made. One is that the lack of reusability unraveled by our analysis is a consequence of independent, unconnected and non-standardised experimental practices. The quality and depth of documentation of Biobricks has been improving over time, but we still see a growing library of unconnected, case-specific constructs with small (or no) reusability. Even if documentation is complete, we lack understanding of what to expect when different, previously unrelated parts are combined for a new construct. An improved theoretical framework is much needed. Secondly, we should also face the possibility of novel ways of thinking about engineering biology. A major difference between SB and its technological counterparts is that synthetic constructs are built to work inside an already functional biological machinery where hardware and software meld. Even if we accept the metaphor of cells as machines (Lazebnik 2002) most of the current SB is intended to add or modify very small parts of an organism that is different in many ways from any existing man-made hardware. Our limited capacity for replacing or connecting multiple parts effectively confines ourselves within the domains not of technology but tinkering.

## Acknowledgments

The authors would like to thank A. Crompton for providing the database of LEGO parts. This work has been supported by an ERC Advanced Grant Number 294294 from the EU seventh framework program (SYNCOM), by the EU seventh framework program (ST-Flow Project), grants of the Botin Foundation, by Banco Santander through its Santander Universities Global Division (SV,RS), by the Spanish Ministry of Economy and Competitiveness, Grant FIS2013-44674-P and FEDER (SV) and by the Santa Fe Institute (RS).

## REFERENCES

1. Milo R, Shen-Orr S, Itzkovitz S et al (2002) Network motifs: simple building blocks of complex networks. Science 298, 824–827.

2. Milo, R., Itzkovitz, S., Kashtan, N at al (2004) Superfamilies of designed and evolved networks. Science 303, 1538–1542.

3. Alon U, (2007) Network motifs: theory and experimental approaches. Nature Rev Genet 8, 450–461.

4. Andrianantonandro E, Basu S, Karig DK and Weiss R (2009) Synthetic biology: new engineering rules for an emerging discipline. Mol Sys Biol msb4100073-E1

5. Arthur B (2009) The Nature of Technology. What it is and How it Evolves. Free Press.

6. Anonymous. Synthetic biology: Beyond divisions. Nature. 2014 May 8;509(7499):151. doi: 10.1038/509151a.

7. Church GM, Elowitz MB, Smolke CD, Voigt CA, Weiss R. Realizing the potential of synthetic biology. Nat Rev Mol Cell Biol. 2014 Apr;15(4):289–94. doi: 10.1038/nrm3767. Epub 2014 Mar 12.

8. Collins J (2012) Synthetic Biology: Bits and pieces come to life. Nature 483: S8?S10

9. Porcar M, Danchin A, de Lorenzo V. Confidence, tolerance, and allowance in biological engineering: the nuts and bolts of living things. Bioessays. 2015 Jan;37(1):95–102. doi: 10.1002/bies.201400091. Epub 2014 Oct 27.

10. Church G and Regis, E (2012). Regenesis. New York: Basic Books. ISBN 0-465-02175-1.

11. de Lorenzo V. Beware of metaphors: chasses and orthogonality in synthetic biology. Bioeng Bugs. 2011 Jan-Feb;2(1):3–7. doi: 10.4161/bbug.2.1.13388.

12. Collins JJ, Maxon M, Ellington A, Fussenegger M, Weiss R, Sauro H. (2014) Synthetic biology: How best to build a cell. Nature 509:155–7.

13. Coulange B (1998) Software reuse. Springer-Verlag, London.

14. Cserer A and Seiringer A (2009) Pictures of synthetic biology. Syst Synth Biol 3:27–35.

15. Dorogovtsev, S.N.; Mendes J.F.F. Evolution of Networks: From Biological Nets to the Internet and WWW; Oxford University Press: Oxford, 2003.

16. Ferrer Cancho R, Janssen C and Solé R (2001) Topology of technology graphs: theall world patterns in electronic circuits. Phys Rev E 64, 046119.

17. Hooper JW and Chester RO (1991) Software reuse. Guidelines and methods. Plenum Press, New York.

18. Maynard-Smith J (2000) The concept of information in biology. Philosophy of Science 67: 177–194.

19. Myers CR (2003) Software systems as complex networks: Structure, function, and evolvability of software collaboration graphs. Phys. Rev. E 68, 046116

20. Newman MEJ (2005) Power laws, Pareto distributions and Zipf’s law. Contemp Phys 46, 323–351.

21. Newman MEJ (2010) Networks: an introduction. Oxford U Press, Oxford UK.

22. Nicholson, DJ. (2013) Organisms ≠ Machines. Studies in History and Philosophy of Biological and Biomedical Sciences 44: 669?678

23. Nurse, P (2008) Life, logic and information. Nature 454: 424?426.

24. Potanin, A., Noble, J., Frean, M., and Biddle, R. (2005) Scale-free geometry in OO programs. Comm. ACM 48, 99–103.

25. Pressman, R. S. (2005) Software engineering: a practitioner’s approach. McGraw-Hill, New York.

26. Purnick P, Weiss R (2009) The second wave of synthetic biology: from modules to systems. Nat Rev Mol Cell Biol 10: 410–422.

27. Solé, R. V., Valverde, S., and Rodriguez-caso, C. (2011) Convergent evolutionary paths in biological and technological networks. Evol Edu Outreach 4: 415–426.

28. Vilanova C, Porcar M. (2014) iGEM 2.0-refoundations for engineering biology. Nat Biotechnol. 32(5), 420–424.

29. Dijkstra, E. W. (1972) The Humble Programmer. Comm. ACM 15(10), 859–866.

30. McIlroy, M.D. (1976) Mass-produced software components. In: Buxton, J.M., Naur, P., Randell, B. (eds.) Software Engineering Concepts and Techniques, 1968 NATO Conference on Software Engineering, Garmisch, Germany, 88–98.

31. IEEE1517-1999 (2004). IEEE1517 Standard for Information Technology - Software Life Cycle Processes - Reuse Processes: 1999, reaffirmed 2004. Software Engineering Standards Committee of the IEEE Computer Society, USA.

32. Garlan, D., Allen, R., and Ockerbloom, J. (1995) Architectural Mismatch: Why Reuse is So Hard. IEEE Software, 17–26.

33. Mili, H., Mili, F., and Mili, A. (1995) Reusing software: uses and research directions. IEEE Trans Software Eng 21, 528–562.

34. Crompton, A. (2012) The Entropy of Lego. Env. Plan. B: Planning and Design 39, 174–182.

35. Solé, R. V., Ferrer-Cancho, R., Montoya, J. M., Valverde, S. (2002) Selection, tinkering, and emergence in complex networks. Complexity 8, 20–33.

36. Teichmann, S., and Babu. M. (2004) Gene regulatory network growth by duplication. Nature Genetics 36, 492–496.

37. Valverde, S., Ferrer-Cancho, R., Solé, R. V. (2002) Scale-free Networks from Optimal Design. Europhys. Lett. 60, 512–517.

38. Valverde, S. and Solé, R. V. (2005) Network motifs in computational graphs: A case study in software architecture. Phys. Rev. E 72, 026107.

39. Valverde, S., Solé, R. V., Bedau M. A., and Packard, N. (2007) Topology and evolution of technology innovation networks. Phys. Rev. E 76, 056118.

40. Yan, K.-K., Fang, G., Bhardwaj, N., Alexander, R. P., and Gerstein, M. (2010) Comparing genomes to computer operating systems in terms of the topology and evolution of their regulatory control networks. Proc Natl Acad Sci USA 107, 9186–9191.

41. Zipf, G. (1936) The Psychobiology of Language. London: Routledge.

42. Walker, S. A. and Davies, P. (2013) The algorithmic origins of life. J. R. Soc. Interface 10, 20120869.

43. Williams, M.R. (1985) A History of Computer Architecture. Prentice-Hall.

44. Baldwin, C. Y. and Clark, K. B. (2000) Design Rules, Volume 1, The Power of Modularity. MIT Press, Cambridge MA.

45. Hey T and Pápay, Y (2015) The computing universe. A journey through a revolution. Cambridge U Press, New York.

46. Tassey, G. (2002) The Economic Impacts of Inadequate Infrastructure for Software Testing. Planning Report 02-3, NIST U.S. Department of Commerce, Technology Administration.

47. Lazebnik V. (2002) Can a biologist fix a radio? - or, what I learned while studying apoptosis. Cancer Cell. 2, 79–82.

